# First occurrence of parasitic isopod (*Braga nasuta*, Schioedte & Meinert, 1881) on ornamental fish in Europe

**DOI:** 10.64898/2025.12.04.692341

**Authors:** Márton Hoitsy, János Gál, Endre Sós, Antal Nagy, Sándor Hornok, Gergő Keve

## Abstract

Parasitic crustaceans represent a significant threat to both freshwater and marine aquaculture, with several families capable of inducing severe diseases with high mortality rates and therefore significant economic losses. Among them, isopods of the genus *Braga* (Cymothoidae) are haematophagous parasites of neotropical fish, causing anaemia, tissue damage, and mortality. *Braga nasuta*, endemic to Brazil, has been reported from various hosts, including ornamental species exported worldwide. During routine examination of imported cardinal tetra (*Paracheirodon axelrodi*, Schultz, 1956), the authors detected that one of the fish harbored an adult male isopod (∼6.21 mm long, 2.66 mm wide) attached anterior to the dorsal fin. Based on morphological identification, the parasite was identified as *Braga nasuta*. No secondary pathogens were observed in affected fish. Given the global trade in ornamental species, the introduction of *B. nasuta* into non-native regions underscores the importance of quarantine and parasite monitoring. While the potential ability of this parasite to establish populations in Europe requires further studies, vigilance is warranted to prevent potential spread and ecological impact. In this study, the authors document the first occurrence of *B. nasuta* in Europe, detected on a cardinal tetra (*P. axelrodi*) imported to a Hungarian public aquarium.

## INTRODUCTION

Parasitic crustacean are well known in fresh and also marine water (Misganaw and Getu, 2016). Some species can cause high losses in marine aquaculture, however the freshwater fish farms are also affected by the obligate aquatic ectoparasites (Abolofia et al., 2017; Haridevamuthu et al., 2024; Noga, 2010). The most relevant freshwater crustacean parasites belong to the Argulidae, Cymothoidae, Ergasilidae and Lernaeidae families (Eiras et al., 2010; Heckmann, 2003).

*Argulus* species can infect ornamental fish also (Mirzaei and Khovand, 2015; Saha and Bandyopadhyay, 2015; Shahraki et al., 2014). The life cycle of the parasite is very similar in Argulidae family. The adult females lay 250-300 eggs to flat surfaces (plants, stones, etc.) in the water. The hatching depends on the water temperature, but usually takes 8-55 days (Fryer, 1982; Molnár and Baska, 2017; Walker et al., 2004). These parasites can cause serious losses, but also can be vectors of several viral diseases (Das et al., 2018; Mikheev et al., 2015; Molnár and Baska, 2017; Molnár and Székely, 1998; Walker et al., 2004).

Compared to the previous parasite the *Ergasilus* spp. are a parasite on the gills of the fish (Abdelhalim et al., 1991). They belong to the Ergasilidae family and mass mortalities can occur in case of heavy infections (Abowei and Ezekiel, 2011; Molnár and Baska, 2017). Only the female specimens are parasitic, which are carrying 20-100 eggs, depending on the species. The pathological signs include gill damage, tissue hyperplasia, lamellae fusions and necrosis. Secondary infections can also manifest on the affected areas (Abdelhalim et al., 1991; Jithendran et al., 2008).

*Lernaea* spp. (anchor worms) also belongs to the group of parasitic crustaceans. They can be found on the external surface of the fish and on the gills. Only the females are parasitic. The anchor worms can cause high mortality in juvenile populations; however, the parasites will cause hemorrhages and inflammation in skin tissues on larger fish. The process can transform into necrosis. Secondary infections can also manifest on the affected area, as bacteria and water molds. These secondary pathogens can induce ulcers and necrosis too (Molnár and Baska, 2017; Noga, 1986).

Parasitic isopods are less common in Europe compared to the previously mentioned crustaceans. Members of this order can infect marine as well as freshwater fish (Nizar et al., 2021; Ravichandran et al., 2009). Representatives of four families are known (Aegidae, Corallanidae, Cymothoidae and Gnathiidae), to have parasitic life stages (Noga, 2010). Larger species can measure up to 50 mm. The species, which belong to the Cymothoidae family are mostly distributed in South America (Thatcher et al., 2003). The genus of *Braga* involves eight species, *Braga amapaensis* (Thatcher, 1996), *Braga bachmanni* (Stadler, 1972), *Braga brasiliensis (*Schioedte & Meinert, 1881), *Braga cichlae* (Schioedte & Meinert, 1881) *Braga cigarra* comb. nov. (Szidat & Schubart, 1960), *Braga nasuta* (Schioedte & Meinert, 1881), *Braga occidentalis* (Boone, 1918) and *Braga patagonica* (Schioedte & Meinert, 1884) (Boone, 1919; Schioedte and Meinert, 1881; Thatcher et al., 2009). These isopods parasitize various neotropical fish. *Braga* species are haematophages and can cause anaemia, skin damage, which can transform into necrosis and ulcers. Some of them live in the abdominal cavity of the fish, while others can be found in the mouth, over the tongue and also reduce opercular movements. All of these can lead to the death of the host (Brandão et al., 2013; Huizing, 1972; Kabata, 1985; Ravichandran et al., 2009; Thatcher, 1996; Thatcher et al., 2009, 2003).

*Braga nasuta* is a parasite of freshwater fish that is endemic in South-America, especially Brazil, where it can be considered widespread. Reports of this parasite originate from various states, such as Amazonas, Bahia and São Paulo (Luque et al., 2013). Furthermore, recent findings of the parasite originate from the State Mato Grosso do Sul from a stream near to the River Paraná, and from the Lower Sao Francisco basin (Alves et al., 2024; Narciso et al., 2019). While the host-preference of the parasite is largely unknown, specimens were retrieved from piraya (*Pygocentrus piraya*, Cuvier, 1819) and from serape tetra (*Megalamphodus eques*, Steindachner, 1882) (Alves et al., 2024; Narciso et al., 2019). In addition, Jesus et al., (2017) found *Braga* sp. they identified as *B. nasuta* on aparaima (*Aparaima gigas*, Schinz, 1822) fingerlings from the Amazonas River. However, based on apparent morphological differences, Narciso et al. (2019) suggested that these specimens may belong to another species of the genus *Braga*. Nevertheless, fish species that can serve as hosts for *B. nasuta* are commonly exported from Brazil as ornamental or aquarium fish (Marková et al., 2020; Narciso et al., 2019). While in their work, Narciso et al. (2019) emphasized that wild-caught fish intended for export may serve as carriers of *B. nasuta*, posing a risk of introduction into importing countries.

The authors present here the first report of *B. nasuta* parasitizing the cardinal tetra fish (*Paracheirodon* axelrodi), sampled in a public aquarium in Hungary.

## MATERIALS AND METHODS

Cardinal tetra (*P. axelrodi*) fish (300 specimens) arrived in the quarantine facility of the public aquarium from a German ornamental fish wholesale. Fish were placed into tank (158×59×59 cm) filled with 500 L water and filtered with Sicce Whale 500 external filtration unit (Sicce S.r.l., Pozzoleone, Italy).

Wet smears (skin scraping, gill biopsy, fecal sample) were taken from the animals which showed clinical symptoms. The samples from the fish were examined by Euromex BioBlue BB.4260 microscope (Euromex Microscopen BV, Duiven, The Netherlands). Pictures (Figures 1&2) were taken with Nikon D7000 (Nikon Corporation, Tokyo, Japan).

**Figure 1:**
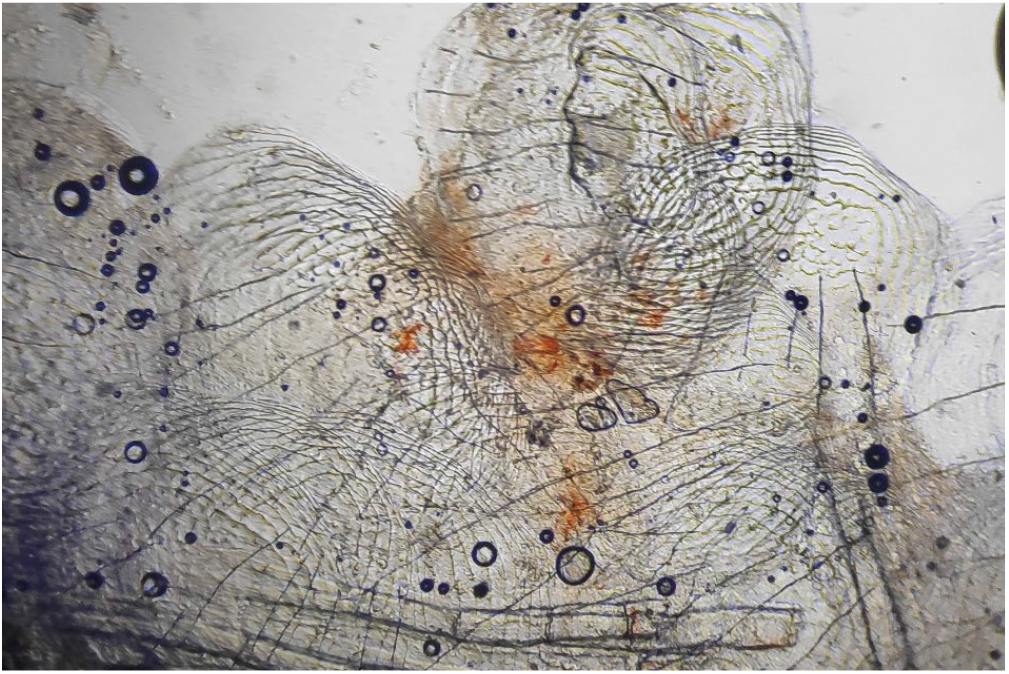
Native wet smear under light microscope from the cardinal tetra (100×)

**Figure 2:**
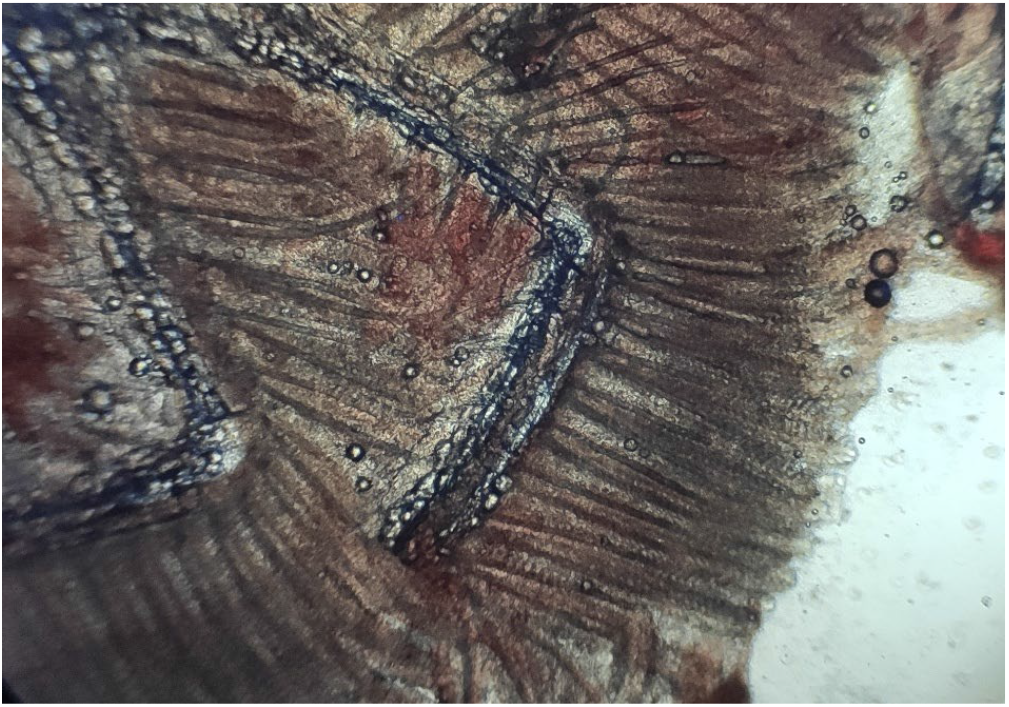
Native gill biopsy under light microscope from the cardinal tetra (100×)

The parasite was removed from the affected fish and stored in 96% ethanol until the morphological identification.

The morphological identification of *B. nasuta* was based on the work of (Narciso et al., 2019).

Photography for Figure (Figures: 3,4,5,6) were made with a stereo microscope and compatible camera Leica DMC4500 and were compiled and measured with LasX (Leica application suite X) program (Leica Microsystems, Wetzlar, Germany). Photography for (Figure 7) were made with light microscope and compatible camera Leica FLEXACAM C1 (LeicaMicrosystems, Wetzlar, Germany) and were compiled with CombineZP program (Hadley, 2010). For measurements, both microscopes were calibrated with the same objective micrometer. Scales were added or adjusted manually.

**Figure 3:**
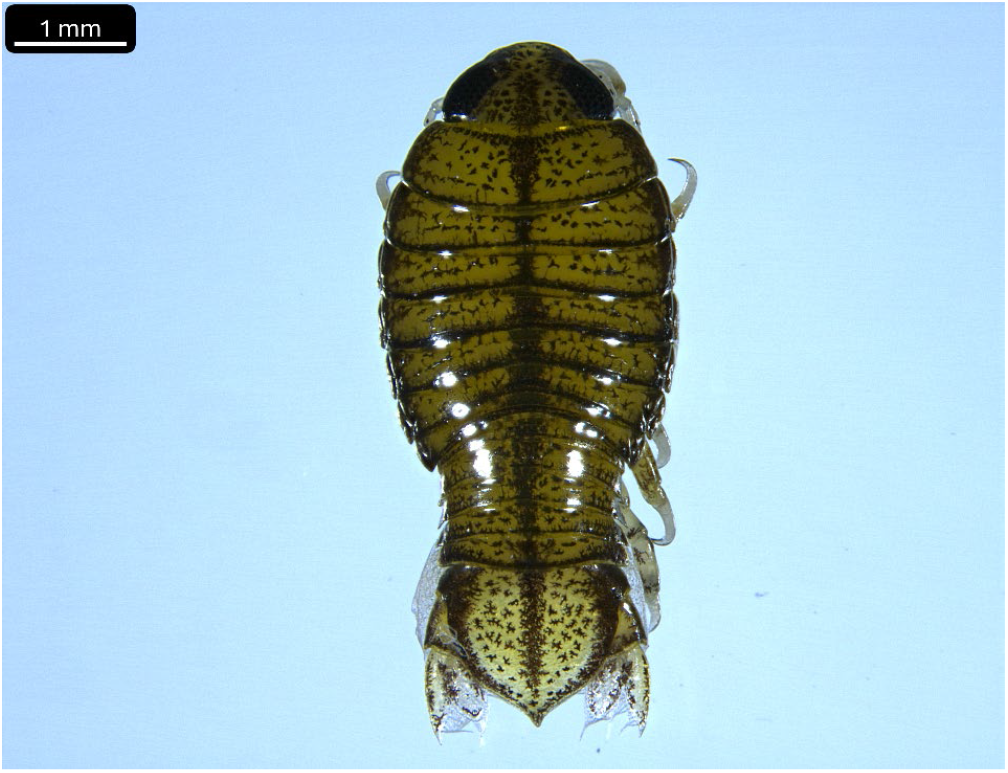
Dorsal aspect of *B. nasuta* (Bar=1 mm)

**Figure 4:**
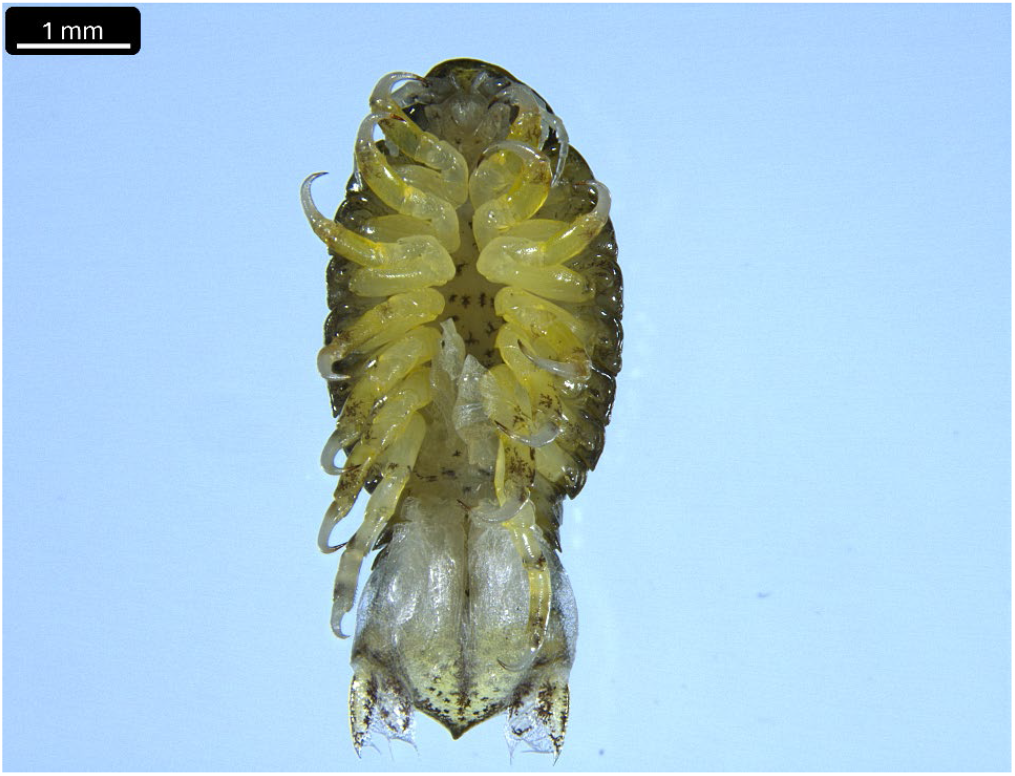
Ventral aspect of *B. nasuta* (Bar=1 mm)

**Figure 5:**
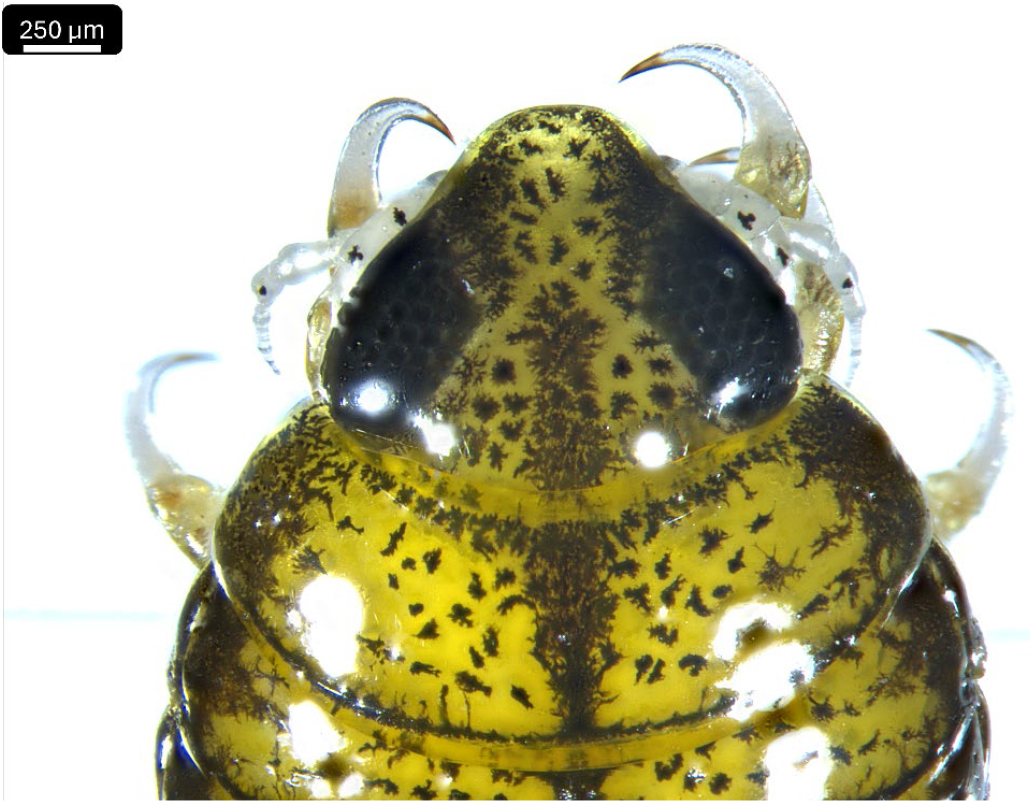
Cephalon of *B. nasuta*, from a dorsal aspect (Bar=250 µm)

**Figure 6:**
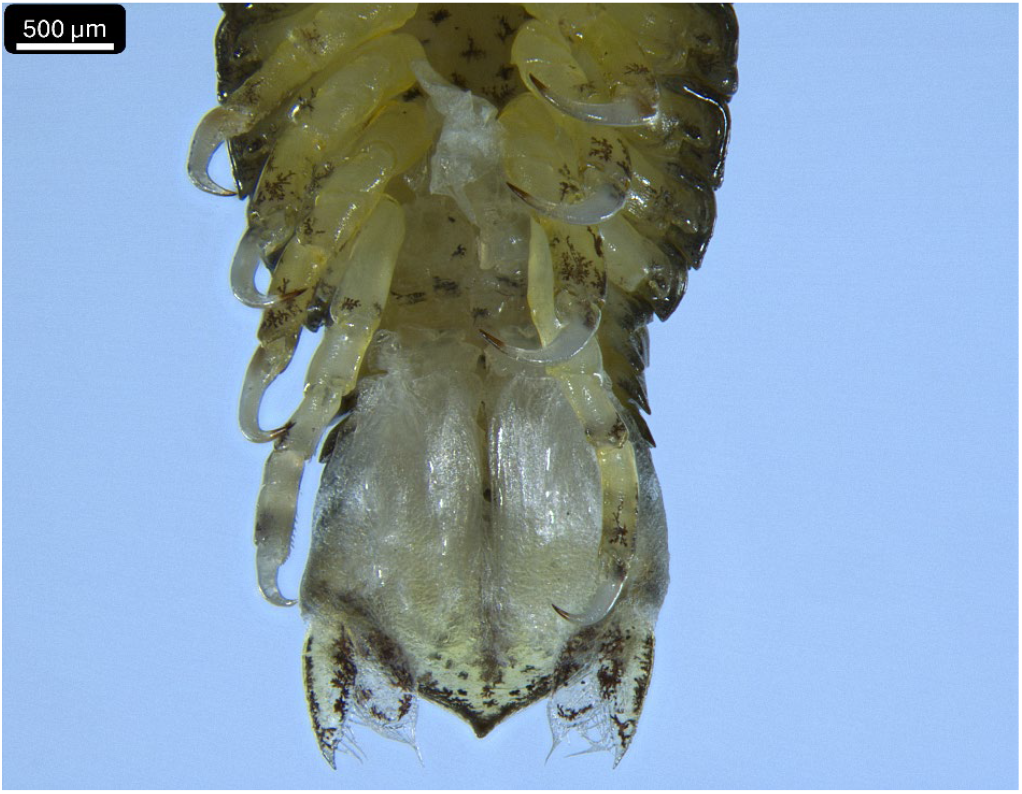
Caudal end of *B. nasuta* from a ventral aspect (Bar=500 µm)

**Figure 7:**
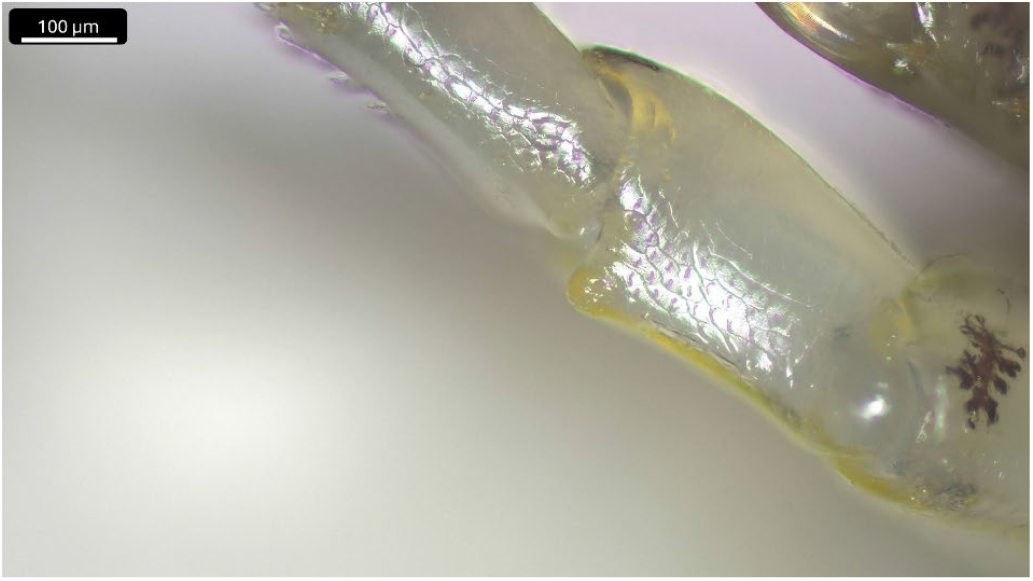
Merus of pereopod 7 with a distal tooth (Bar=100 µm)

## RESULTS

Three hundred specimens of cardinal tetra (*P. innesi*) were examined for external parasites.

The affected cardinal tetra fish showed irregular swimming behavior. Whitish, grayish patches (1-3 mm diameter) were visible on skin of the animals. Sporadic mortalities were in the quarantine tank. The examined fish were anaemic. The skin scrapes from the fish did not show any ciliate or fungal pathogens. Gill biopsies were clear from parasitic species. The structure of gill filaments looked normal, but pale on the native samples under the microscope (Figures 1&2). The fecal samples from the intestine were normal, parasites were not observable.

Parasitic isopod (∼6.21 mm long, 2.66 mm wide) was visible on the external surface of the cardinal tetra front of the dorsal fin. Just one fish specimen hosted an adult *male* isopod (prevalence: 0.33%).

Following the previously mentioned work (Narciso et al, 2019), the identification of the isopod was based on the following features: (A) body symmetrical, longer than wide (∼6.21 mm long, 2.66 mm wide) (Figures 3&4)); (B) cephalon not immersed in pereonite 1 (Figure 5); (C) all seven pairs of pereopods prehensile and provided with stout claw-like dactyls (Fig. 4); (D) Pleopods multilamilate (Figures 4&6); (E) pleon slightly immersed in pereonite 7 (Figure 3); (F) cephalon truncated anteriorly (Figure 5); (G) pereopod 7 with a dactyl shorter and weaker than others (Figure 6); (H) merus of pereopod 7 with a distal tooth near the articulation with carpus (Figure 7); (I) pleotelson triangular, ending in a small and sharp tip (Figure 6); and (J) uropod extending beyond the apex of pleotelson and with exopod slightly longer than endopod (Figure 6).

Based on the morphological analysis, the single specimen was identified as *Braga nasuta*.

## DISCUSSION

Fish parasites can spread worldwide easily with international transport. The new ornamental fish can be affected with parasites which can infect the local fish species to (Lymbery et al., 2014; Williams et al., 2013). Co-introduced parasites are arriving with the host and can spread in the natural environment also. The crustacean parasites are infesting local species, such as the Japanese fish louse (*Argulus japonicus*, Thiele, 1900) on the Hungarian common carp (*Cyprinus carpio*, Linnaeus, 1758). Moreover, this invasive crustacean that can already be considered to be widespread in Europe, may also be able to establish new, geographically and molecularly different sub-species in the continent (Keve et al., 2025). Furthermore, ornamental crustaceans themselves (such as such as *Neocaridina davidi)* can potentially act as vehicles of non-native parasites during international transport (Hoitsy et al., 2023).

Based on the morphological analysis, the single specimen was identified as *B. nasuta*, although our specimen was somewhat larger than the specimen described in the work of Narciso et al, (2019), ∼6.21 mm long, 2.66 mm wide vs 5.6 mm long and 2.6 mm wide. A small difference was also observed in the length/width ratios: 2.33 vs 2.15. This said, our specimen was considerably smaller than those described in the work of Jesus et al., (2017): 14.06 ± 2.3 mm long by 6.46 ± 1.2 mm wide. Note, that in their work, Narciso et al. (2019) compared their results to those of Jesus et al. (2017). According to the former, the specimens reported by Jesus et al. (2017) likely belong to a different species of the same genus (*Braga*). Despite this, it is noteworthy that sheer size differences were not mentioned in their differential diagnosis. In their work Alves et al. (2024) reported a single male specimen of *B. nasuta* collected from its natural habitat. According to their depiction (calculated from: Alves et al., 2019: Figure 3) this specimen was approximately 7.07 mm long and 3.05 mm wide (length/width ratio: 2.31). While these small differences may have arisen from methodological differences, it is worth mentioning that in the case of other crustacean parasites such as *A. japonicus*, it is known that males (and females) can be considerably different in size as per different life stages (Keve et al., 2025).

Our morphological identification, as well as the previous suspicion of Narciso et al (2019) is further strengthened by the fact that the German ornamental fish wholesale had claimed that they are importing wild caught fish species from Brazil, where *B. nasuta* is native.

On our specimen, we observed pigmentations on the antennulae as well as on the antennae (Fig.3). Similar pigmentations are visible on photography in the work of Narciso et al. (2019): Figure 2/A, although these are not depicted on the drawings of Alves et al. (2024): Figures 3/C, D.

Although the appearance of *B. nasuta* in countries that import ornamental fish from Brazil was foreshadowed by Narciso et al, (2019), to the best of our knowledge this is the first time this parasite was identified in Europe. The potential ability of *B. nasuta* to establish wild populations in European countries require further studies, however we consider this scenario unlikely given the difference in climates between Brazil: mainly tropical and sub-tropical (Alvares et al., 2013) and Europe: mainly mediterranean, oceanic and continental (Deliège and Nicolay, 2016). However, this cannot be excluded based on the emergence of invasive fish species in Hungary (continental climate), some of which originate from South- and Central-America (Takács et al., 2017). Nevertheless, our result highlights the importance of monitoring imported animals for the presence of parasites.

## Acknowledgements

The authors would like to thank the colleagues of the Department of Exotic Animal and Wildlife Medicine and Clinic and Department of Parasitology and Zoology at the University of Veterinary Medicine, Budapest, for their assistance, and the collaboration of the Tropicarium Oceanarium Kft. (Public Aquarium) for providing the samples, and for making the publication of this case.

## Authors contributions

MH: Manuscript writing, sample collection, data curation, conceptualization, photography. JG, ES, AN, SH: Critical revision of the manuscript, result validation. GK: Manuscript writing, morphological identification, photography, data curation, conceptualization

## Funding

Financial support was provided by the Office for Supported Research Groups, Hungarian Research Network (HUN-REN), Hungary (project no. 1500107).

## Ethics approval and consent to participate

Not applicable.

## Consent for publication

Not applicable.

## Competing interests

The authors declare no competing interests

## Notes

### Competing Interest Statement

The authors have declared no competing interest.

